# Deep Learning Explains the Biology of Branched Glycans from Single-Cell Sequencing Data

**DOI:** 10.1101/2022.06.27.497708

**Authors:** Rui Qin, Lara K. Mahal, Daniel Bojar

## Abstract

Glycosylation is ubiquitous and often dysregulated in disease. However, the regulation and functional significance of various types of glycosylation at cellular levels is hard to unravel experimentally. Multi-omics, single-cell measurements such as SUGAR-seq, which quantifies transcriptomes and cell surface glycans, facilitate addressing this issue. Using SUGAR-seq data, we pioneered a deep learning model to predict the glycan phenotypes of cells (mouse T lymphocytes) from transcripts, with the example of predicting β1,6GlcNAc-branching across T cell subtypes (test set F1 score: 0.9351). Model interpretation via SHAP (SHapley Additive exPlanations) identified highly predictive genes, in part known to impact (i) branched glycan levels and (ii) the biology of branched glycans. These genes included physiologically relevant low-abundance genes that were not captured by conventional differential expression analysis. Our work shows that interpretable deep learning models are promising for uncovering novel functions and regulatory mechanisms of glycans from integrated transcriptomic and glycomic datasets.

## Introduction

Glycosylation is a ubiquitous post-translational modification of proteins. Approximately half of all proteins are glycosylated (Apweiler et al., 1999), influencing their physical properties and biological activities (A. Varki et al., 2017; Varki, 2017). Cells utilize glycosylation to control their function and fate. For example in CD8^+^ T lymphocytes, β1,6-branched glycans of CD8^+^ T cell surface proteins are upregulated following T cell activation to prevent overstimulation. Downregulation of sialylated core 1 *O*-linked glycans of CD8^+^ T cell surface proteins induces apoptosis in the absence of activation, maintaining T cell homeostasis (Van Dyken et al., 2007; Priatel et al., 2000; Smith et al., 2018). Changes in glycosylation have been mechanistically implicated in cancer, infectious diseases, autoimmune diseases, metabolic disorders, and developmental defects (Demus et al., 2021; Ng and Freeze, 2018; Pinho and Reis, 2015; Qin and Mahal, 2021; Reily et al., 2019; Sun et al., 2016; Theodoratou et al., 2014; Vosseller et al., 2002).

Despite many observational studies reporting glycosylation changes in diseases, much remains unknown about the origin of these changes. Glycan biosynthesis is orchestrated in a non-templated manner by hundreds of enzymes encoded by “glycogenes”, including glycosyltransferases (add sugar residues), glycosidases (remove sugar residues), sugar modifying enzymes (synthesize phosphorylated, sulfated, or acetylated glycans), enzymes of sugar metabolism pathways, and sugar transporters (A. Varki et al., 2017; Neelamegham and Mahal, 2016). Regulation of glycosylation includes transcriptional and post-transcriptional control of glycogenes, substrate availability, and intracellular trafficking of enzymes (Neelamegham and Mahal, 2016). Therefore, identifying factors driving observed changes in certain glycan structures can be a formidable challenge.

The functional significance of disease-associated glycosylation changes can also be difficult to ascertain, due to the multi-modal influences of glycans. Glycans can influence protein structure, interactions between proteins and receptors, recognition by carbohydrate-binding lectins, resistance to endocytosis and protease degradation, etc (A. Varki et al., 2017; Johannes et al., 2018; Mimura et al., 2018). One glycan feature can thus have multiple effects. For example, increased α2,3-sialylation of cancer cells contributes to cancer progression and metastasis through mechanisms including (i) immune system evasion by interacting with α2,3-sialic acid-binding, immunosuppressive Siglec receptors (e.g., Siglec-9), (ii) increasing metastatic potential via selectins (specific for α2,3-sialic acid-containing sialyl Lewis x and sialyl Lewis a antigens) displayed on circulating cells, and (iii) promoting angiogenesis and the epithelial-to-mesenchymal transition (EMT) process (Dobie and Skropeta, 2021; Natoni et al., 2016; Pietrobono and Stecca, 2021; Rodriguez et al., 2021).

Analyzing multi-omic data has recently emerged to identify glycosylation-related mechanisms in pathogenesis. Combining transcriptomic and glycomic data identified factors driving melanoma metastasis, pancreatic cancer, and HIV persistence (Agrawal et al., 2017; Colomb et al., 2020; Kurz et al., 2021). Agrawal et al. examined RNA-seq datasets of melanoma and found increased transcript levels of an enzyme synthesizing core fucosylated glycans in metastasized melanoma, matching melanoma glycosylation profiles. They also reported more transcripts of transcription factors that directly upregulate the expression of the core fucose-synthesizing enzyme (Agrawal et al., 2017).

State-of-the-art machine learning (ML) approaches, such as deep learning (DL) algorithms, are increasingly used to map transcriptomes onto phenotypic differences, such as tissue types, cancer stages and grades, drug responses, and disease outcomes, to understand and explain phenotypic outcomes on a molecular systems level (Hanczar et al., 2020; Jia et al., 2021; Smith et al., 2020; Yap et al., 2021). Model explanation methods such as Integrated Gradients (Dincer et al., 2018), LRP (layer-wise relevance propagation) (Bach et al., 2015), LIME (Local Interpretable Model-agnostic Explanations) (Ribeiro et al., 2016), DeepLIFT (Deep Learning Important FeaTures) (Shrikumar et al., 2017), and SHAP (SHapley Additive exPlanations) facilitated using DL models to shed light on biological questions. SHAP unifies and improves precedent methods (Lundberg and Lee, 2017). It assigns a “SHAP value” to each input feature, reflecting its impact on the expected output of a DL model for an input example. Using SHAP to identify biologically relevant transcripts contributing to phenotypic differences has only been explored recently (Huang et al., 2022; Withnell et al., 2021; Yap et al., 2021). Traditionally, these were identified by differential expression analysis (DEA), which is (i) prone to information loss due to arbitrariness regarding p-value and fold change thresholds (Bui et al., 2020; Yang et al., 2019) and (ii) biased towards highly expressed genes (Oshlack and Wakefield, 2009). Yap et al. found predictive genes identified by SHAP that were not identified by DEA (Yap et al., 2021), indicating that SHAP can detect subtle but important differences.

We hypothesized that we could develop a DL model to differentiate glycosylation states of single cells and use model interpretation approaches such as SHAP to provide meaningful biological insights into glycan biosynthesis and function (Figure 1A). To the best of our knowledge, DL algorithms using transcripts as inputs have not been employed to study differential glycosylation before. Our model is based on data from a new technology (SUGAR-seq) that simultaneously measures the transcriptome and glycosylation in single cells (Kearney et al., 2021). Technologies such as SUGAR-seq capture the microheterogeneity at single cell levels, which is inaccessible by bulk omics yet can be valuable for mechanistic interpretation, and also provide high volume, matched multi-omics data that enable ML modelling. Specifically, the SUGAR-seq data comprise single-cell transcriptomes of mouse T lymphocytes and the abundance data of surface β1,6-branched glycans on these cells. Using the transcriptomic data as input to predict the binarized glycosylation phenotype of the cells, our model achieved an average F1 score of 0.9351 on the test set, with comparable performance across cell types. It also outperformed alternative approaches, including Random Forest and Gradient Boosting. SHAP analysis identified highly predictive genes (“SHAP genes”) that are involved in the biology of branched glycans in various ways, including their biosynthesis. SHAP genes were also enriched in immunosuppression pathways, matching known functions of branched glycans in T cells. Importantly, SHAP genes included low-abundance genes, such as glycogenes, which were not captured by DEA. Our work shows that explainable DL models are a promising tool for uncovering novel functions and regulatory mechanisms of glycans from paired single-cell transcriptomic and glycomic data.

**Figure 1.**
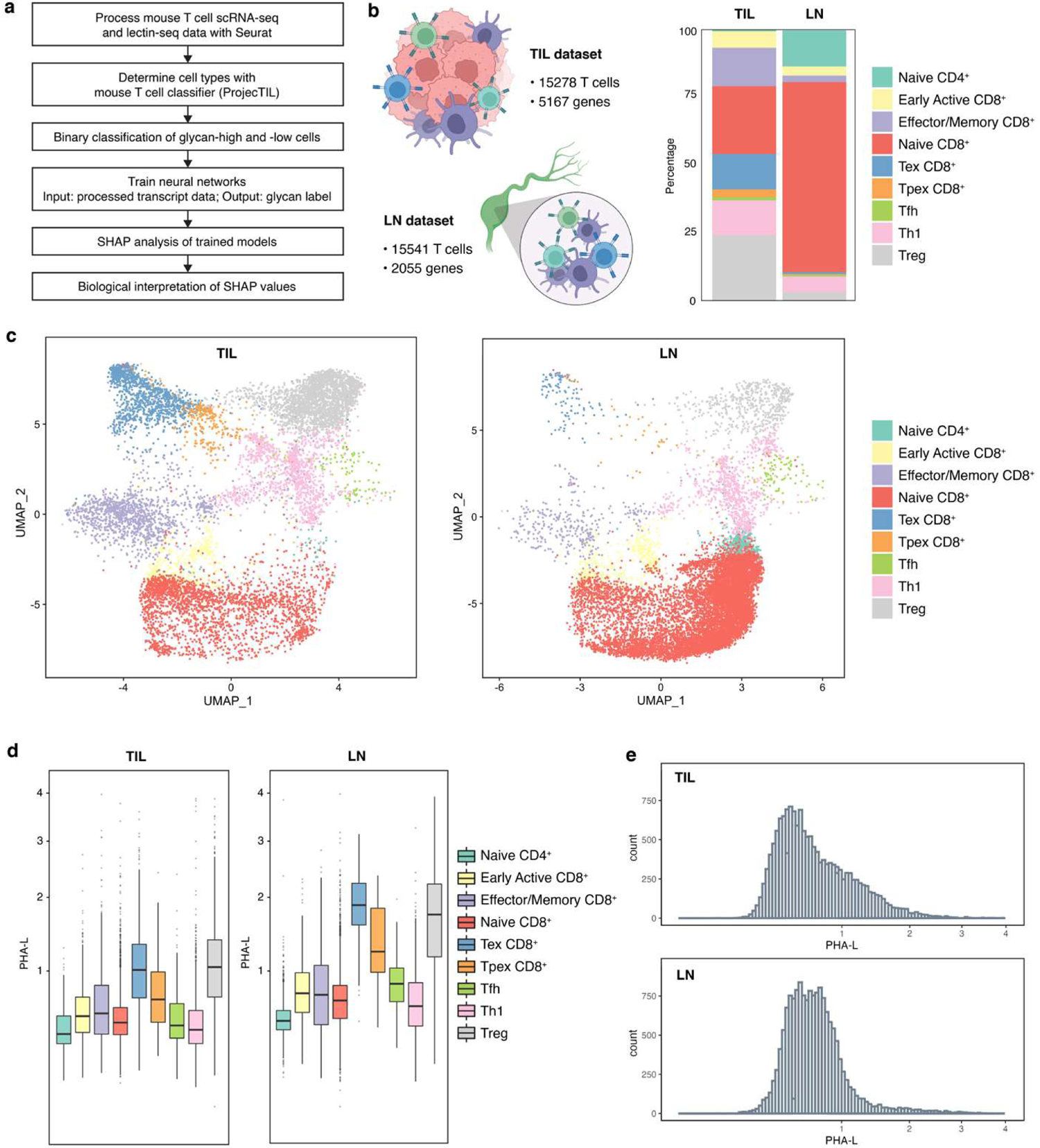
Different mouse T cells show distinct cell surface glycosylation patterns based on single cell RNA- and lectin-seq. (a) Graphical summary of workflow. (b) Composition of the TIL and LN datasets. Left: numbers of cells and genes in each processed dataset. Right: cell type composition as percentages in each dataset. (c) UMAP clustering of cells in each dataset. (d) Boxplots of processed PHA-L data (β1,6-branched glycan abundance) by cell type in each dataset. (e) Histograms of processed PHA-L data in each dataset. Cell type annotations: Tex, terminally exhausted T cell; Tpex, precursor exhausted T cell. Tfh, follicular helper T cell; Th1, T helper 1 cell; Treg, regulatory T cell.

## Results

### Differential Expression of Surface β1,6-branched Glycans in Mouse T Cells

Extracting mechanistic insights into the role of glycans within the context of a cell would be greatly aided by paired single-cell data combining multiple systems biology modalities. We thus used recently publicly available single-cell RNA- and lectin-seq data (Kearney et al., 2021), which included two datasets corresponding to mouse (i) tumor-infiltrating T lymphocytes (TIL) and (ii) lymph node T lymphocytes (LN) (Figure 1B). *Phaseolus vulgaris* leucoagglutinin (PHA-L) was used for lectin-seq. PHA-L is highly specific for branched *N*-glycans with β1,6-GlcNAc linkage (“β1,6-branched glycans”), a glycoform implicated in tumor progression, tumor metastasis, and immune cell development and functional regulation (Bojar et al., 2022; Demetriou et al., 2001; Granovsky et al., 2000; Morgan et al., 2004; Mortales et al., 2020). PHA-L binding is a proxy for the activity of alpha-1,6-mannosylglycoprotein 6-beta-*N*-acetylglucosaminyltransferase A (MGAT5), the enzyme synthesizing the β1,6-GlcNAc linkage. Based on transcriptomes, T cells were categorized into 9 major subtypes, including naïve/naïve-like CD4^+^ T cells, naïve/naïve-like CD8^+^ T cells (may include central memory T cells), early active CD8^+^ T cells, effector memory CD8^+^ T cells, terminally exhausted CD8^+^ T cells (Tex), precursor exhausted T cells (Tpex), regulatory T cells (Treg), T helper 1 cells (Th1), and follicular helper T cells (Tfh) (Figure 1B; Figure 1C). We validated this classification via marker gene expression (Figure S1). Additionally, TIL composition is highly heterogeneous, comprising activated and regulatory cells. Conversely, the LN pool mainly contained resting T cells (Figure 1B; Figure 1C). Observed compositions of T cells isolated from different sites are consistent with previous studies (Kumar et al., 2018; Szabo et al., 2019).

Surface glycosylation varies by cell type (Agrawal et al., 2014; Holst et al., 2016; Tao et al., 2008). We observed clear and reproducible differences in surface expression of β1,6-branched glycans across T cell subtypes in both datasets (Figure 1D). Treg and Tex consistently exhibited the highest expression of branched glycans, while naïve CD4^+^, naïve CD8^+^, and Th1 exhibited the lowest. Similarly, previous studies reported greater PHA-L binding to stimulated T cells than to their naïve counterparts (Cabral et al., 2017; Smith et al., 2018). Glycan expression was also more variable in TIL, consistent with its more diverse composition (Figure 1E). As surface branched glycans seemed to distinguish T cells of different characteristics and functions, we hypothesized that we could use DL to uncover the biology behind this differential glycosylation.

### Training A Deep Learning Model to Predict Glycan Phenotypes from the Transcriptome

We set out to use DL to model surface glycosylation from transcriptome-wide gene expression data. We assigned binary labels to cells based on PHA-L data, corresponding to high and low ends of reads (top 25% “PHA-L^high^”, bottom 25% “PHA-L^low^”). This robustly separated biologically distinct populations and enabled subsequent analyses. It also transformed our task into binary classification, with gene expression values as input, and the probabilities for the positive phenotype (PHA-L^high^) of the corresponding cells as output.

We developed a neural network classifier comprising four hidden layers (Figure 2A), using data from either the TIL or the LN dataset. In the TIL hold-out test set, this classifier achieved a 92.17% prediction accuracy for the PHA-L^high^ phenotype and 95.01% for the PHA-L^low^ phenotype (AUC: 0.9359, F1 score: 0.9351, Table 1; Figure 2B). In the LN dataset, F1 score was lower, yet still exceeded 0.91. Prediction accuracies for both phenotypes remained greater than 90% (Table 1). This slight decrease in performance was most likely due to less variance in the LN data. Robustness of the classifier was also indicated by non-overlapping distributions for the predicted probabilities of PHA-L^high^ (Figure 2C). We further observed comparable performance across different cell subtypes, despite the inherent imbalance in T cell compositions (Figure 2D).

**Figure 2.**
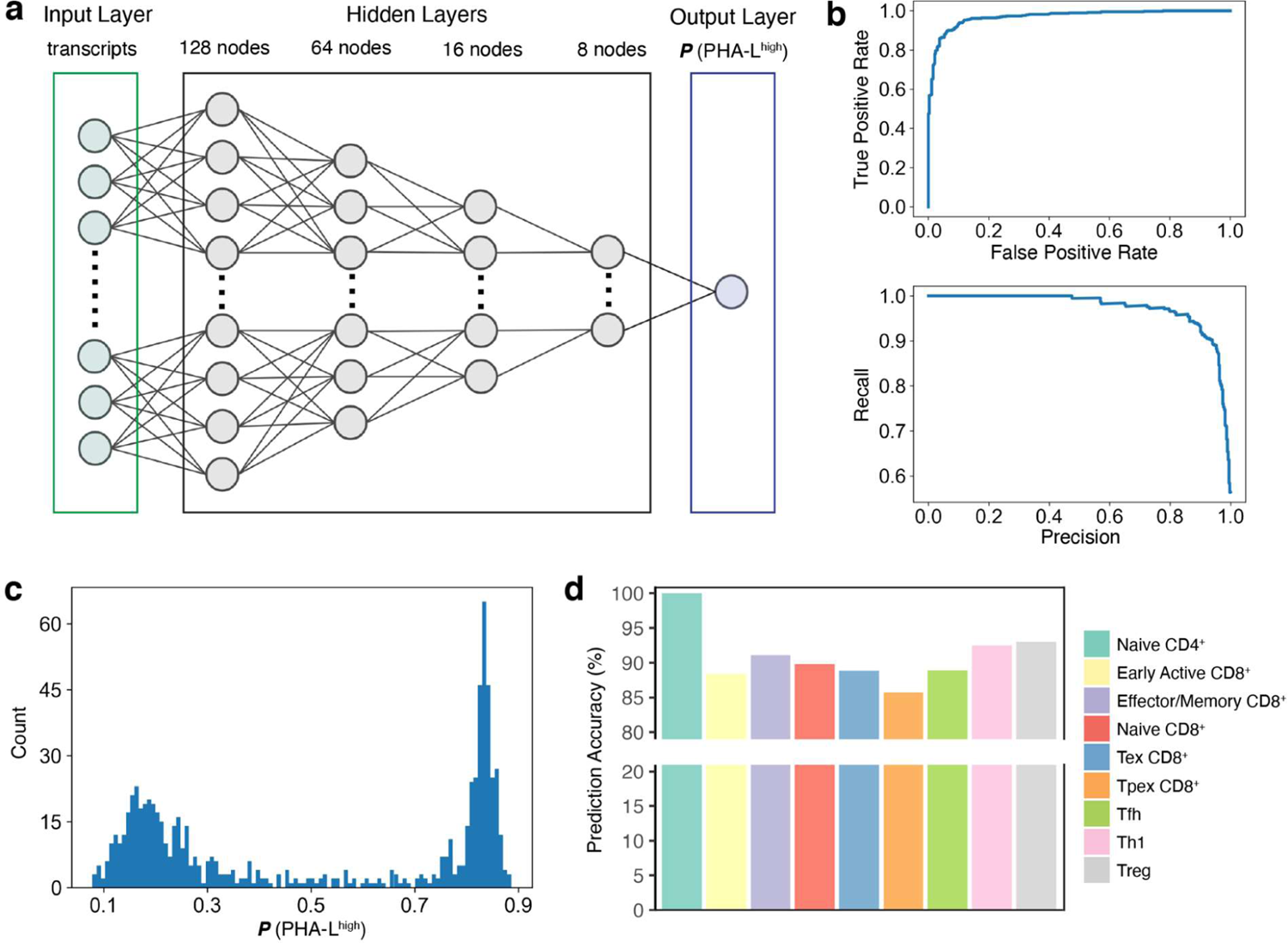
The deep learning model trained on TIL dataset is highly accurate in predicting glycan classes. **(a)** Graphical description of the neural network structure. **(b)** ROC curve (upper) and precision-recall curve (lower) of the model using the test set data. **(c)** Histogram of model output (probability for PHA-L^high^ class) using the test set data; **(d)** Prediction accuracies by cell types of the test set.

**Table 1.** Performance metrics of the model.

|  | TIL |  |  | LN |  |  |
| --- | --- | --- | --- | --- | --- | --- |
|  | Training set | Validation set | Test set | Training set | Validation set | Test set |
| Prediction Accuracy for PHA-L<sup>high</sup> | 95.21% | 90.42% | 92.17% | 94.93% | 93.15% | 90.26% |
| Prediction Accuracy for PHA-L<sup>low</sup> | 97.85% | 91.69% | 95.01% | 92.26% | 93.94% | 92.49% |
| Average Cross Entropy Loss | 0.2031 | 0.2802 | 0.2485 | 0.3088 | 0.3323 | 0.3346 |
| ROC Curve AUC<br>(Positive class: PHA-L<sup>high</sup>) | 0.9653 | 0.9106 | 0.9359 | 0.9360 | 0.9204 | 0.9137 |
| F1 score<br>(Positive class: PHA-L<sup>high</sup>) | 0.9649 | 0.9102 | 0.9351 | 0.9371 | 0.9217 | 0.9131 |

<sup>a</sup> TIL: tumor-infiltrating T lymphocyte; LN: lymph node T lymphocyte.
<sup>b</sup> ROC Curve AUC: the area-under-the-curve of the receiver operating characteristic curve

Using the TIL data, we trained three alternative models (Convolutional Neural Network (CNN), Random Forest, AdaBoost) on the same task and compared their performance to the abovementioned neural network model (Table 2). For all investigated metrics, our neural network outcompeted alternative models. Among the alternative models, Random Forest had the highest prediction accuracies (90.34% for PHA-L^high^, 95.01% for PHA-L^low^) and F1 score (0.9251). In contrast, the CNN model had the lowest prediction accuracies (89.82% for PHA-L^high^, 91.60% for PHA-L^low^) and F1 score (0.9065), although the cross-entropy loss was comparable to the standard network. We thus used the neural network model for all subsequent analyses.

**Table 2.** Comparison of performance metrics of different models to predict glycan classes in the TIL dataset.

|  | Standard Neural Network |  | Convolutional Neural Network |  | AdaBoost |  | Random Forest |  |
| --- | --- | --- | --- | --- | --- | --- | --- | --- |
|  | Training set | Test set | Training set | Test set | Training set | Test set | Training set | Test set |
| Prediction Accuracy for<br>PHA-L<sup>high</sup> | 94.21% | 92.17% | 91.18% | 89.82% | 90.09% | 90.86% | 91.07% | 90.34% |
| Prediction Accuracy for<br>PHA-L<sup>low</sup> | 97.85% | 95.01% | 92.93% | 91.60% | 93.03% | 92.91% | 96.53% | 95.01% |
| Average Cross Entropy Loss | 0.2031 | 0.2485 | 0.2035 | 0.2499 | 0.6690 | 0.6556 | 0.2891 | 0.2901 |
| ROC Curve AUC<br>(Positive class: PHA-L<sup>high</sup>) | 0.9653 | 0.9359 | 0.9205 | 0.9071 | 0.9156 | 0.9188 | 0.9380 | 0.9268 |
| F1 Score<br>(Positive class: PHA-L<sup>high</sup>) | 0.9469 | 0.9351 | 0.9200 | 0.9065 | 0.9145 | 0.9182 | 0.9346 | 0.9251 |

<sup>a</sup> ROC Curve AUC: the area-under-the-curve of the receiver operating characteristic curve

### Identification of Highly Predictive Genes Using SHAP

Next, we used SHAP to identify genes important for prediction. For each input sample (cell), the SHAP algorithm calculates a “SHAP value” for each feature (gene) that is reflective of the impact of this feature on the expected model output for this input (Lundberg and Lee, 2017). SHAP values can be positive or negative, corresponding to additive or subtractive effects on model output. The median absolute SHAP value is commonly used to assess global feature importance.

We averaged SHAP values derived from three identical models with comparable performance, separately trained using different seeds for random splitting of datasets. Then we calculated median absolute SHAP values for all genes (Table S1), resulting in gene rankings. For both the TIL and the LN model, only a small portion of genes were highly important (Figure 3A). We chose genes corresponding to the top 10% SHAP rankings (“SHAP genes”), yielding 516 genes for TIL and 205 genes for LN (see Figure 3B (TIL) and Figure S2 (LN) for examples). For most SHAP genes (e.g., *Lag3*, *Ccl5*, *Foxp3*, Figure 3B), higher expression tended to favor a PHA-L^high^ prediction, whereas the opposite was true for a smaller subset (e.g., *Gm42418*, *Malat1*, *Klf2*, Figure 3B). As glycogenes directly control glycan biosynthesis (Neelamegham and Mahal, 2016), we also examined the top 30 glycogenes (Figure 3C), although not all were among the top 10% SHAP genes.

**Figure 3.**
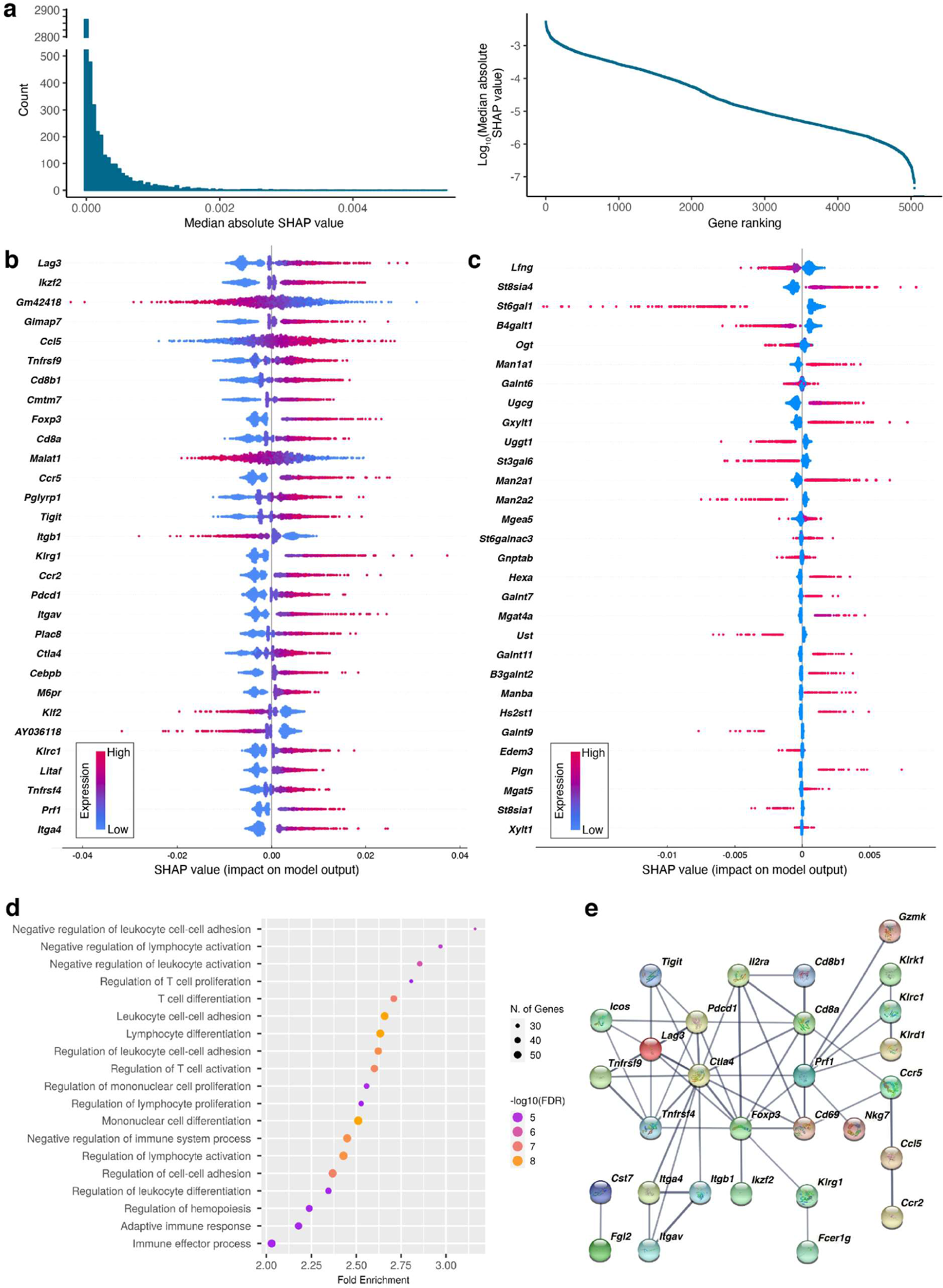
Model interpretation identifies genes important for predicting TIL cell surface glycosylation. (a) Histogram (left) and scatter plot (right) presentation of the median absolute SHAP values for all genes. (b) SHAP values of top 30 genes ranked by median absolute SHAP value. (c) SHAP values of top 30 glycogenes ranked by median absolute SHAP value. (d) Gene Ontology pathway enrichment analysis of using the SHAP genes. (e) STRING protein interaction network analysis of the top 10%SHAP genes. Only high confidence (strong evidence) interactions are shown, and thicker edges denote higher confidence. Genes/proteins without high confidence interactions with any other genes/proteins are not displayed.

Next, we performed pathway enrichment analysis on the SHAP genes. For both the TIL and LN model, the most enriched pathways (biological processes) were involved in negative regulation of T cell activity and T cell differentiation (TIL: Figure 3D; LN: Figure S3), matching known roles of β1,6-branched glycans in T cells discussed further below. STRING protein interaction analysis also showed the top 10% SHAP genes to be highly interconnected via functional enrichment, co-expression, or direct interaction, arguing for concerted biology (Figure 3E).

Next, we investigated the variation of SHAP genes with T cell subtypes, generating cell type-specific lists of SHAP genes (Table S2). As shown in Figure 4A, the majority of highly predictive genes were shared across cell subtypes. For example, *Ctla4* (cytotoxic T-lymphocyte-associated protein 4) was consistently highly ranked in all cell subtypes (highest rank-naïve CD4^+^: 13 or top 0.25%; lowest rank-Tex: 165 or top 3.19%). *Ctla4* is a Treg marker, with research primarily focusing on its roles in Treg (Rowshanravan et al., 2018; Sobhani et al., 2021; Zheng et al., 2013). Our analysis showed that *Ctla4* expression was predictive of glycan phenotype and might influence biological pathways mediated by branched glycans in T cells beyond Treg.

**Figure 4.**
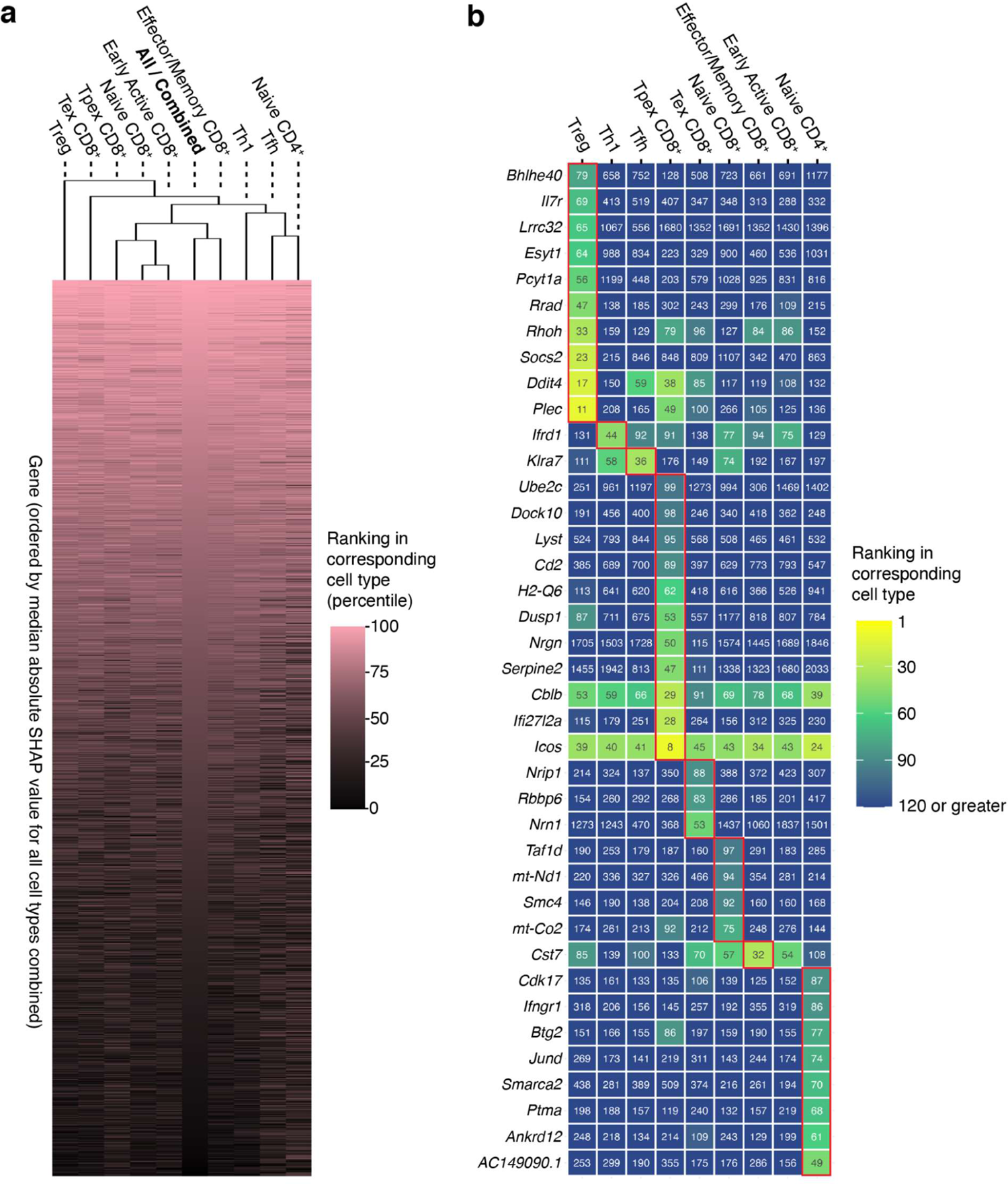
Most of the highly predictive genes are shared across cell types in the TIL dataset. (a) Heatmap of the percentage rankings (by median absolute SHAP value) of genes in each cell type compared to all cell types combined. Cell types are clustered by euclidean distance. (b) Genes uniquely high ranked (median absolute SHAP value among top 2%) in each cell type. Values in tiles are the rankings of genes (by median absolute SHAP value) in the corresponding cell types. The highest rankings of genes among all cell types are boxed in red.

Nonetheless, we identified some genes that were highly predictive of glycan phenotype only in certain T cell subtypes. A gene is considered specifically important to a T cell subtype if its ranking percentile in this subtype is (i) among the top 2% and (ii) greater than the average ranking percentile in other subtypes by at least 1.25 standard deviations. This identified gene subsets for each cell subtype (Figure 4B, TIL). Treg, naïve CD4, and Tpex exhibited the highest numbers of type-specific highly predictive genes. Some genes play unique biological roles in their corresponding subtypes and are associated with the biology of branched glycans. For example, our cell-type specific comparison ranked *Lrrc32* (leucine rich repeat containing 32; also known as glycoprotein-A repetitions predominant, *Garp*) as particularly high in Treg. *Lrrc32* controls the expression of latent TGF-β (transforming growth factor β) in Treg, and TGF-β signaling is suppressed in *Mgat5* knockout mice (Lehmkuhl et al., 2021; Tran et al., 2009; Zhang et al., 2021). We speculate that *Lrrc32* may mediate this loss of TGF-β signaling, explaining its importance for prediction in this context. *Jund* (transcription factor JunD) and *Ifngr1* (interferon gamma receptor 1), highly ranked in naïve CD4^+^ T cells, are important for the activation and differentiation of naïve CD4^+^ cells, usually followed by upregulating β1,6-branched glycans (Afkarian et al., 2002; Meixner et al., 2004; Morgan et al., 2004). *Jund* and *Ifngr1* expression may reflect differential levels of activation in naïve CD4^+^ cells, which could correlate with branched glycan expression. As most other cell subtype-specific genes lack well-characterized roles in the corresponding subtypes of T cells, their biological associations with branched glycans are unclear and constitute opportunities for future research.

### SHAP Genes Explain the Biology of β1,6-branched Glycans

One of our major goals was to see whether the most predictive SHAP genes, identified by this data-driven approach, were implicated in the biology of branched glycans. A highly predictive gene can be biologically associated with β1,6-branched glycans in several ways: (i) encoding a cell surface protein bearing this glycan; (ii) regulating the expression of MGAT5, the enzyme biosynthesizing the β1,6-branch of *N*-glycans; (iii) being itself regulated by MGAT5/β1,6-branched glycan levels; (iv) being functionally synergistic with β1,6-branched glycans; (v) influencing the biosynthesis of β1,6-branched glycans through, e.g., changing substrate availability. Indeed, many highly predictive SHAP genes (i.e., SHAP ranking in top 2%), were already implicated in β1,6-branched glycan biology in at least one of these ways (Figure 5; Figure 6), arguing for high biological relevance of SHAP genes. We detail our findings below.

**Figure 5.**
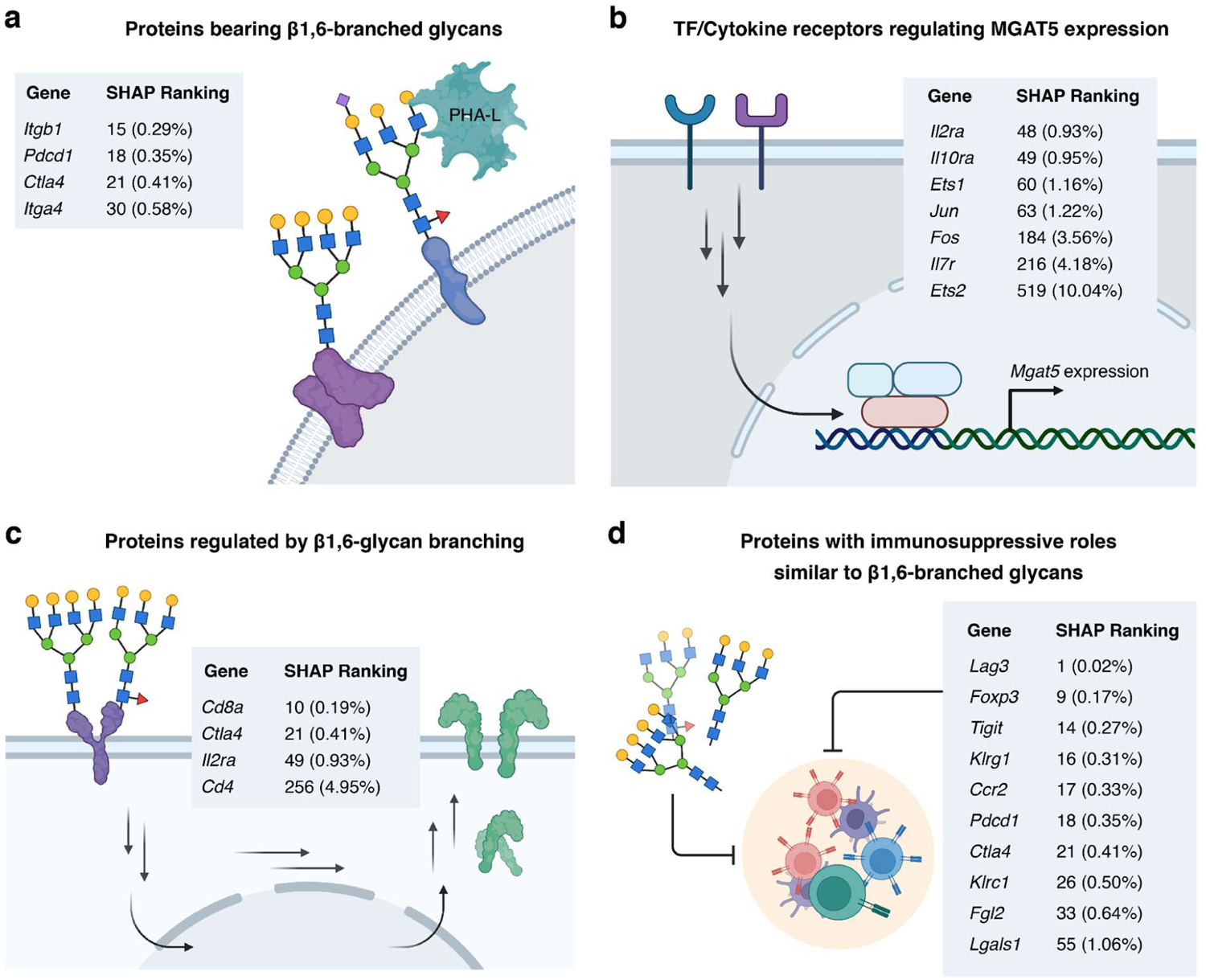
Predictive genes identified by SHAP tend to be involved in the biology of MGAT5/β1,6-branched glycans. These genes encode proteins that: **(a)** bear PHA-L binding, β1,6-branched N-glycans that can be important to their protein functions; **(b)** regulate the expression of MGAT5/β1,6-branched glycans; **(c)** are regulated by β1,6-glycan branching; **(d)** have immunosuppressive functions that may be synergistic with β1,6-branched N-glycans, which are also immunosuppressive. Gene names, their rankings (by median absolute SHAP value) and relative rankings (ranking/number of all genes × 100%, indicated in parentheses) are shown.

**Figure 6.**
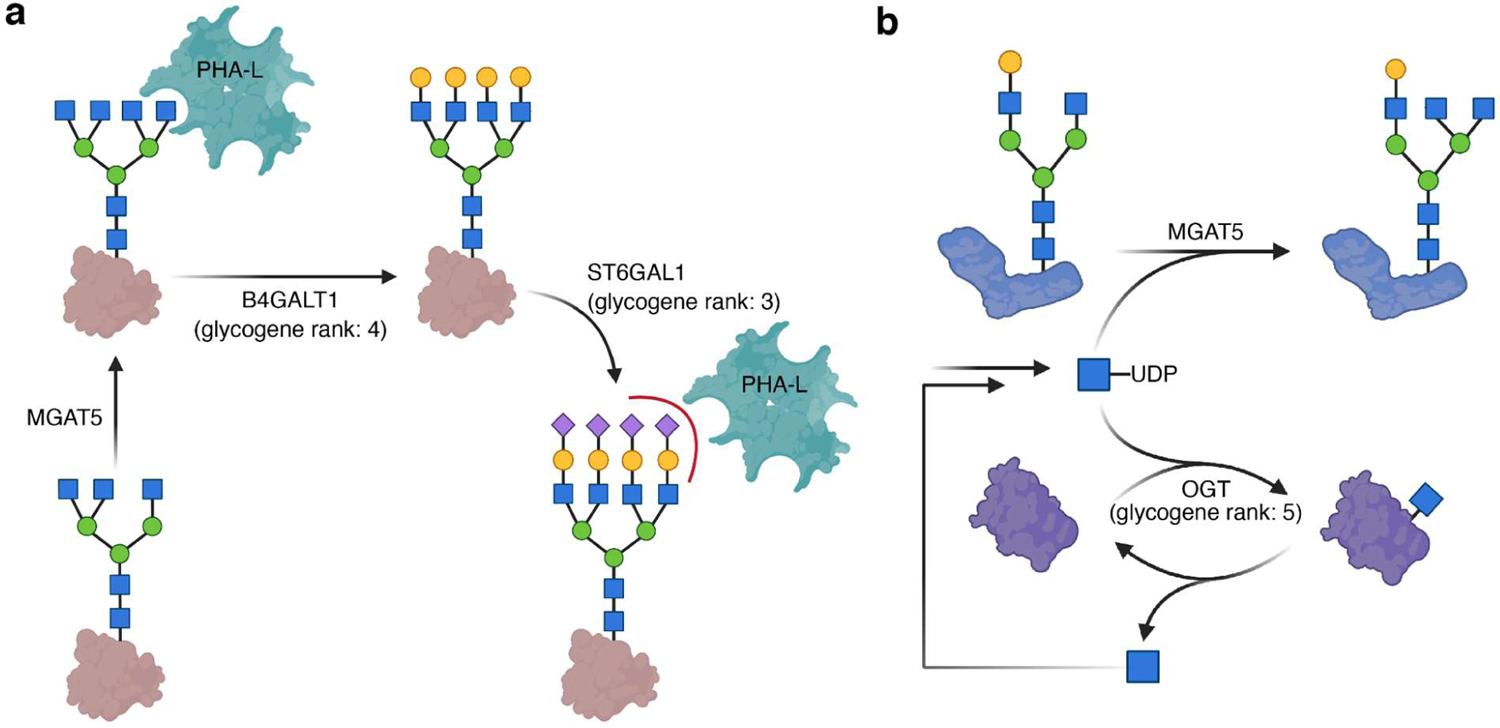
SHAP analysis identifies glycogenes that impact β1,6-branched N-glycan levels or PHA-L binding. (a) Partial biosynthetic route of ɑ2,6-sialylated, β1,6-branched glycans and the involvement of B4GALT1 and ST6GAL1 identified by SHAP analysis in this process. ɑ2,6-sialylation abrogates PHA-L binding to branched N-glycans. **(b)** Interplay between UDP-GlcNAc, O-GlcNAcylation and N-glycan branching, and the involvement of OGT in this process. UDP: uracil diphosphate group.

We primarily focused on interpreting SHAP genes from the TIL dataset because the dataset is more representative of a general T cell population, and the corresponding model exhibited better prediction performance. First, we investigated whether TIL SHAP genes encode T cell surface proteins bearing β1,6-branched glycans, as some glycoproteins are more prone to be modified with this glycoform (Breen, 2002; Li et al., 2013; Wu et al., 2019). Out of the top 50 SHAP genes, *Pdcd1* (programmed cell death protein 1, PD-1), *Ctla4*, *Itgb1* (integrin beta 1), and *Itga4* (integrin alpha 4) can be modified with β1,6-branched glycans (Figure 5A) (Liu et al., 2020; Przybyło et al., 2007; Zhu et al., 2014). Except for *Itgb1*, these genes had SHAP values increasing with expression, as expected. Others either lack experimental glycosylation data or encode non-membrane proteins. Identifying proteins modified with β1,6-branched glycans among the most predictive genes may inform future investigations on whether other SHAP genes also encode proteins with this glycoform.

Next, we investigated SHAP genes regulating MGAT5 expression. Only three transcription factors seem to directly upregulate MGAT5 expression: ETS-1, ETS-2, and AP-1 (dimer of c-Jun and c-Fos) (Chen et al., 1998; Ko et al., 1999; Wang et al., 2018). In our SHAP analysis, *Ets1*, *Jun*, and *Fos* were all ranked among the top 2% genes (Figure 5B), coinciding with their importance in regulating MGAT5 expression. The ranking of *Ets2* was lower, potentially due to lower expression (Figure S4), but still bordering the top 10% range (Figure 5B). Intriguingly, SHAP values of *Ets1* negatively correlated with expression, potentially due to cell-dependent regulation. Indeed, RNA-seq data showed increased *Mgat5* transcripts in T cells of *Ets1^-/-^* mouse (Kim et al., 2018), suggesting ETS-1-mediated downregulation in T cells. Beyond transcription factors, we also investigated more upstream cytokine regulators. Three cytokines, interleukin-2 (IL-2), interleukin-7 (IL-7), and interleukin-10 (IL-10), upregulate β1,6-branched glycans in T cells (Grigorian et al., 2012; Smith et al., 2018). Correspondingly, *Il2ra*, *Il10ra*, *Il2rg*, and *Il7r*, encoding components of the membrane receptors of IL-2, IL-10, IL-2/IL-7, and IL-7, are highly ranked at top 0.93%, 0.95%, 3.79%, and 4.18%, respectively (Figure 5B). They were also the top four ranked genes out of the 29 interleukin receptor genes here, indicating their strong association with glycan branching. In aggregate, all transcription factors and cytokine receptors regulating MGAT5/branched glycan levels were highly predictive SHAP genes, showing the potency of SHAP analysis to reveal the mechanisms behind the regulation of glycosylation.

Dysregulated T cell-mediated immunity has been identified in *Mgat5*^-/-^ animals (Demetriou et al., 2001; Lee et al., 2007; Silva et al., 2020). Therefore, we hypothesized some SHAP genes could be regulated by MGAT5/branched glycans. Deleting *Mgat5* decreases surface expression of CTLA-4, CD8 alpha coreceptor, and CD4 coreceptor in T cells, while upregulating *Mgat5*/surface β1,6-branched glycans resulted in increased T cell surface expression of IL-2 receptor (Araujo et al., 2017; Lau et al., 2007; Zhou et al., 2014). These studies argue MGAT5 upregulate these proteins was by rescuing them from endocytosis. However, whether the mRNA levels of these proteins also changed was not investigated. In our analysis, the corresponding genes, *Ctla4*, *Cd8a*, *Cd4*, and *Il2ra*, were highly ranked at 0.41%, 0.19%, 4.95% and 0.93% (Figure 5C). This suggests that regulation of these proteins by glycan branching may involve both mRNA expression and endocytosis.

Next, we sought to interpret genes controlling glycan biosynthesis. We identified four glycogenes, *Lfng*, *St8sia4*, *St6gal1*, and *B4galt1*, among the top 10% SHAP rankings. ST6GAL1 (Beta-galactoside alpha-2,6-sialyltransferase 1) is the predominant enzyme adding terminal α2,6-linked sialic acid to galactose residues in glycans. While this step occurs downstream of *N*-glycan branching (Petit et al., 2010), α2,6-sialylation blocks the binding of β1,6-branched glycans to PHA-L and endogenous receptors such as galectins (Figure 6A) (Bojar et al., 2022; Stowell et al., 2008; Zhuo et al., 2008). Thus, SHAP values of *St6gal1* were negatively correlated with expression (Figure 3C). The same was found for *B4galt1* (beta-1,4-galactosyltransferase 1), the primary enzyme transferring β1,4-galactose to glycans (Bydlinski et al., 2018). B4GALT1 is indispensable for efficient α2,6-sialylation (Figure 6A), potentially explaining the decreasing SHAP values upon *B4galt1* expression (Khoder-Agha et al., 2019; Nguyen et al., 2021).

Expression of the fifth-ranking glycogene, *Ogt* (total rank: 10.06%), decreased with SHAP values (Figure 3C). OGT transfers GlcNAc onto intracellular proteins. The activated sugar donor UDP-GlcNAc is the shared substrate of MGAT5 and OGT. Supplementing GlcNAc, converted to UDP-GlcNAc in cells, upregulates both *O*-GlcNAcylation and β1,6-glycan branching (Araujo et al., 2017; Taylor et al., 2009). Conversely, a drug-induced decrease in cellular UDP-GlcNAc levels affected both *O*-GlcNAcylation and β1,6-glycan branching (Ricciardiello et al., 2018). OGT thus influences substrate availability for MGAT5, altering abundances of β1,6-branched glycans (**Figure 6B**). A new study showed that shRNA knockdown of OGT indeed increased β1,6-glycan branching (Song et al., 2022).

Having identified biological associations between highly ranked SHAP genes and branched glycans, we asked whether we could systematically identify functional roles of glycan branching in T cells. We based this on previous observations that cells utilize both proteins and protein glycosylation to fulfill their function. For example, fucosylated glycans and sialyl Lewis x on effector T cell surfaces facilitate homing to tumor sites, as do mechanisms mediated by chemokine receptors and integrins (Alatrash et al., 2019; Sackstein et al., 2017). Work over the past two decades has shown β1,6-glycan branching to play an immunosuppressive role in activated T cells. T cell receptor (TCR) activation upregulates MGAT5, yielding more β1,6-branched glycans on T cells. This promotes binding of the multimeric galectin-3 to T cell surface glycoproteins, forming a localized lattice that prevents T cell-activating protein-protein interactions (e.g., TCR-CD8 interaction) (Demetriou et al., 2001; Lau et al., 2007; Morgan et al., 2004). Mice deficient in *Mgat5* developed autoimmune disease due to dampened negative regulation of T cell activities (Grigorian and Demetriou, 2011; Silva et al., 2020). In Treg, surface β1,6-branched glycans were positively correlated with immunosuppressive marker expression and the suppressive potency of Treg (Cabral et al., 2017). Ye et al. identified *Mgat5* as one of the four hits in a CRISPR screen for targets that enhance T cell-based cancer therapy (Ye et al., 2019). Correspondingly, functional enrichment of SHAP genes showed enrichment in negative regulation of T cell activation (Figure 3D). Well-established suppressive immune checkpoint receptors were within the top 50 genes, such as *Lag3*, *Ctla4*, *Pdcd1*, and *Tigit* (Figure 5D). We also found other highly ranked SHAP genes known for predominantly immunosuppressive roles in T cells, including: (i) genes of the killer cell lectin-like receptor family such as *Klrc1* (0.31%) and *Klrg1* (0.50%) (Huot et al., 2021; Li et al., 2016); (ii) chemokines and chemokine receptors such as *Ccl5* (0.10%), *Ccr5* (0.21%), and *Ccr2* (0.33%) (Aldinucci and Casagrande, 2018; Matsuo et al., 2021; Tu et al., 2020; Zeng et al., 2022); (iii) other genes such as *Fgl2* (0.64%) and *Lgals1* (1.06%) (Corapi et al., 2018; Hou et al., 2021) (Figure 5D). Overall, SHAP genes had substantial overlap with genes known to be implicated in the biology of branched glycans.

Finally, we examined the SHAP genes of the LN dataset (Table S1; Figure S2). More than half (116 genes) of the 205 SHAP genes of the LN dataset were also found among the 16 SHAP genes of the TIL dataset. Examples include *Cd8b1* (0.04% in LN, 0.14% in TIL), *Ikzf2* (0.83% in LN, 0.04% in TIL), *Gm42418* (0.13% in LN, 0.06% in TIL), and *Ets1* (4.33% in LN, 1.16% in TIL). Notably, the negative correlation between the transcript abundance and SHAP values of *Ets1* was also seen in the LN dataset. Shared high-ranking genes could be associated with the biology of β1,6-branched glycan in similar ways as discussed above. Some genes had substantially different rankings, which was anticipated since the two datasets had different compositions of T cell subtypes. For example, the highest-ranking gene, *Lag3* (lymphocyte activation gene 3), in TIL was ranked at 746 (36.30%) in LN, due to the minimal expression of *Lag3* in resting T cells comprising most of the LN dataset (Grosso et al., 2007). The roles of β1,6-branched glycans in resting T cells are much less understood. *Mgat5*^-/-^ mice displayed a lowered T cell activation threshold (Demetriou et al., 2001), suggesting that β1,6-branched glycans are also immunosuppressive in resting T cells, by desensitizing them to stimulus. Therefore, we again hypothesized that the SHAP genes of the LN dataset are primarily immunosuppressive, synergizing with β1,6-branched glycans. In line with this, SHAP genes of the LN dataset are functionally enriched in the pathway of negative regulation of T cell activity (Figure S3). The top 50 SHAP genes of the LN dataset also included immunosuppressive genes, such as *Cxcr6*, *Socs1*, *Cd7*, and *Klrd1* (Heesch et al., 2014; Pace et al., 2000; Sempowski et al., 2004; Sheu et al., 2005; Takahashi et al., 2011). *Stab1*, known for its anti-immunosuppression activity in T cells, was ranked at 31 and had SHAP values decreasing with expression (Beyer et al., 2011; Nüssing et al., 2019; Stephen et al., 2017). One study showed that IL-7 treatment reduced β1,6-glycan branching in resting T cells, in contrast to its effect in activated T cells (Mkhikian et al., 2011). Aligning with this, the SHAP values of *Il7r* (rank 2.29%) anti-correlated with mRNA expression in the LN dataset in contrast to the correlation observed in the TIL dataset, underscoring the context-aware nature of our approach.

### SHAP Genes Include Low Abundance Genes That Are Not Captured by Differential Expression Analysis

SHAP identifies gene subsets that were not captured by differential expression analysis (DEA) (Yap et al., 2021). We wondered whether this could also be observed here, and whether any SHAP-exclusive genes were biologically relevant. Using the TIL data, we performed DEA between PHA-L^high^ and PHA-L^low^ populations. Applying a false discovery rate threshold of 0.05, we identified 381 differentially expressed genes (7.4% of all genes; Table S3), slightly less than the number of SHAP genes (516 genes or 10% of all genes). 267 genes were present in both DEA and SHAP genes, with many shared top-ranking genes (e.g., *Lag3*, *Tigit*, *Malat1*).

DEA is known for biasing towards highly expressed genes (Bui et al., 2020; Yang et al., 2019). Among the DEA genes, 18 (4.72%) were highly-transcribed ribosomal genes (Petibon et al., 2020; Zhao et al., 2018), with two even among the top 50 DEA genes. In contrast, only 8 (1.55%) ribosomal genes were found in the SHAP genes, none of which were in the top 50. Other high-abundance housekeeping genes were also found only in DEA genes, such as *Gapdh* (rank 23), *Pgk1* (rank 60), and *Actb2* (rank 71). These observations suggest that SHAP is less prone to biasing towards high-abundance genes. Pointedly, transcription factors *Jun* and *Ets2*, known to upregulate β1,6-glycan branching as discussed above, did not appear in DEA genes. Moreover, glycogenes in general were absent from DEA genes. Overall, our results indicate SHAP to be more powerful than the more conventional DEA in identifying low-abundance genes that are biologically important for phenotypic glycosylation differences.

## Discussion

In contrast with the profusion of genomic, transcriptomic, and proteomic data, matching glycomic data remain scarce (Qin and Mahal, 2021). Consequently, glycosylation research has long been lacking integrated multi-omics analysis and has not benefited greatly from rapidly evolving computational tools for mapping molecular interaction networks in health and disease (Kellman and Lewis, 2021). State-of-the-art artificial intelligence technologies, used widely for drug response prediction or regulatory molecule identification (Adam et al., 2020; Kim et al., 2021; Zheng et al., 2020), remain rarely used in glycosylation studies. Single-cell RNA- and lectin-seq technologies such as SUGAR-seq have started to provide new opportunities to use DL for studying differential glycosylation, as demonstrated here.

It should be noted that the feasibility of predicting glycan features from single-cell transcriptomics data cannot be considered a foregone conclusion. Most genes that would seem relevant from a domain perspective (glycosyltransferases, sugar transporters, etc.) are typically lowly expressed and frequently absent from single-cell transcriptomics data (Nairn et al., 2008; Qiu, 2020). Nonetheless, we show that it is feasible to use the sparse nature of single-cell data to predict a glycan feature with high accuracy using a neural network model. Models predicting “cross-omics”, in this case from the realm of the transcriptome towards the glycome, will be important in the future to aid in multi-omics integration and are particularly relevant in the context of glycans, as they are technically outside the central dogma of molecular biology. Our results here, however, yet again demonstrate that glycans can be re-integrated with the rest of the central dogma, by using transcriptional information to successfully predict parts of glycan expression.

Using SHAP, we further showed that, next to predicting glycan feature abundance, our model can also be used to extract, at scale, compelling biological associations between the transcriptome and the glycome. Tellingly, the genes most important for glycosylation phenotype prediction significantly overlapped with genes involved in the biology of β1,6-branched glycans. A direct explanation for the high predictivity of some SHAP genes could be that they encode membrane proteins bearing β1,6-branched glycans. Although we only identified four genes with experimental evidence for β1,6-branching glycosylation among the top 50 TIL SHAP genes (*Pdcd1*, *Ctla4*, *Itgb1*, *Itga4*), branched glycans are involved in functions for two of them: In persisting exhausted T cells, PD-1 interacts with galectin-9, which binds to branched glycans displayed on PD-1, to inhibit galectin-9-mediated cell death (Yang et al., 2021). Integrin beta 1 is an essential component of a series of integrin complexes that are critical mediators of T cell adhesion and signaling, and its activities are regulated by β1,6-branching of glycans (Bellis, 2004; Jankowska et al., 2018). We expect that more highly-predictive SHAP genes will be discovered to bear β1,6-branched glycans in immune cells, influencing protein function. *Lag3*, a heavily glycosylated immune checkpoint receptor implicated in many diseases (Graydon et al., 2021), was the most predictive gene in the TIL dataset. Although the glycosylation profile of this protein has not yet been determined, LAG3 binds galectin-3 in T cells, an immune system lectin with similar binding specificity as PHA-L (Demetriou et al., 2001; Kouo et al., 2015). This indicates that LAG3 function may be regulated by β1,6-branched glycans. Thus, our DL-based approach can inform future investigations on whether other SHAP genes encode proteins with this glycoform and whether it might impact protein function.

While most SHAP genes do not encode surface glycoproteins, they can still be associated with the biology of β1,6-branched glycans in multiple ways, as discussed above. We note that all known transcription factor- and cytokine-regulators of MGAT5/β1,6-branched glycans, as well as some glycogenes impacting β1,6-branched glycan synthesis, were among the top ranked SHAP genes. This emphasizes the potential of a DL-based approach to identify regulatory mechanisms of glycosylation. Examining the top ranked SHAP genes also yielded new candidates for potential regulators of β1,6-branched glycans. The chemokine receptor CCR2 (C-C chemokine receptor type 2; rank 0.33%), when bound to its ligand CCL2 (C-C motif chemokine 2), upregulates the transcription factor AP-1 that increases MGAT5 expression (Fei et al., 2021; Lin et al., 2012). In turn, AP-1 also enhances CCL2 expression (Deng et al., 2013; Fei et al., 2021; Novoszel et al., 2021). Therefore, the CCL2/CCR2 axis may be an alternative mechanism to regulate MGAT5/β1,6-branched glycans in immune cells. In line with the immunosuppressive role of β1,6-branched glycans, CCL2 secreted by cancer cells contributes to an immunosuppressive tumor microenvironment, and blockade of CCR2 in mice improved the efficacy of immune checkpoint therapy (Matsuo et al., 2021; Tu et al., 2020), suggesting a direct biomedically relevant role of this glycan feature.

Our work showed that (i) glycosylation phenotypes can be modeled by neural networks from transcriptomic data, (ii) biological processes linked to post-translational modifications such as glycosylation can be deciphered by DL model-explaining methods such as SHAP, and (iii) this combined approach may facilitate the discovery of new regulators of branched glycans and downstream effectors of branched glycan-mediated pathways. We note that this single model essentially recapitulates decades of experimental work on this aspect of biology, including generated hypotheses for future work. While we report largely overlapping regulatory associations in our set of immune cell types, future work could also compare these results with the regulation and function of β1,6-branched glycans in other cell types or other species, to develop a global understanding of the diverse roles of this glycan feature.

We envision a pipeline in which this explainable DL approach is used to analyze data generated by SUGAR-seq or similar technologies, to decode the biology of less well-studied glycoforms (e.g., high-mannose glycans, sulfated glycans, I-branched glycans) and their importance in disease (Chuzel et al., 2021; Dimitroff, 2019; de Leoz et al., 2011; Loke et al., 2016; Sun et al., 2022). For this, future work needs to expand the capabilities of the associated lectin-seq, similar to the already reported Glycan-seq technology (Oinam et al., 2021). We also anticipate the integration of more data modalities besides transcriptomes. In RNA-seq, measurement of glycogenes and other low-abundance genes is less accurate, potentially skewing the interpretation of the importance of these genes (Tarazona et al., 2011). Adding proteomic or miRNA data (important regulators of low abundance genes) to the DL model input may provide a more accurate account of the regulation of glycosylation mediated by low-abundance genes (Schmiedel et al., 2015; Thu and Mahal, 2020). We envision that this combination of systems biology and artificial intelligence will provide fresh insights into the complex, interleaved biosynthesis and functional role of glycans in various biological contexts.

## Acknowledgements

This work was funded by a Branco Weiss Fellowship – Society in Science awarded to D.B., by the Knut and Alice Wallenberg Foundation, and the University of Gothenburg, Sweden. Some graphical contents were created with Biorender.com.

## Author contributions

Conceptualization, D.B.; Methodology, R.Q. and D.B.; Formal Analysis, R.Q. and D.B.; Investigation, R.Q. and D.B.; Writing – Original Draft, R.Q. and D.B.; Writing – Review & Editing, R.Q., L.K.M., and D.B.; Supervision, L.K.M. and D.B.

## Data availability

The matrices of processed RNA and PHA-L reads, and the R and python codes for single-cell data processing, model training and SHAP analysis have been uploaded to https://github.com/BojarLab/scGlycomics_b16_branching.

## Methods

### Dataset, Data Processing, and T Cell Subtype Classification

10X sequencing data of mouse tumor infiltrating T lymphocytes (TIL) and mouse lymph node T lymphocytes (LN) were downloaded from Gene Expression Omnibus (GSE166325, GSE166326). Single-cell data processing was performed with the Seurat package (version 4.0.6) in R (Hao et al., 2021). For each dataset, downloaded raw data were initialized with the Read10X function of Seurat. Doublets were removed with the HTODemux function of Seurat. Next, RNA counts were normalized and scaled with the SCTransform function of Seurat, in which the mitochondrial genes, ribosomal genes, and cell cycle phase scores (computed with the CellCycleScoring function of Seurat) were regressed out. The default residual variance cut-off (1.3) of SCTransform was used and the low variance genes were removed. PHA-L reads were processed with the Normalize function of Seurat, using the centered log ratio transformation method.

T cell subtype classification was performed with the multi-dataset reference atlas-based mouse T cell classifier ProjecTIL (version 2.0.0) in R (Andreatta et al., 2021). ProjecTIL categorized the remaining T cells into 9 major subtypes, including naive and naïve-like CD4^+^ T cells (naïve CD4^+^), naïve and naïve-like CD8^+^ T cells (naïve CD8^+^), early active CD8^+^ T cells (early active CD8^+^), effector memory CD8^+^ T cells (effector memory CD8^+^), terminally exhausted CD8^+^ T cells (Tex), precursor exhausted T cells (Tpex), regulatory T cells (Treg), T helper 1 cells (Th1), and follicular helper T cells (Tfh). Cell subtype was confirmed by comparing marker gene expressions (Figure S1).

### Model Training

To generate input data from the exported matrix of transcript and PHA-L reads, cells of PHA-L reads within the upper quartile range (top 25%) were assigned the label 1 (PHA-L^high^), and cells of PHA-L reads within the lower quartile range (bottom 25%) were assigned the label 0 (PHA-L^low^). Other cells were removed. Input data was randomly split into a training set (72%), validation set (18%), and test set (10%). All models (standard neural network, convolutional neural network, Random Forest, and AdaBoost) were trained using the same training, validation, and test set in Python 3.8.

For the standard neural network, models were trained using the PyTorch framework. Models were trained in mini-batches of size 128. For the best performing model, the neural network consisted of 4 hidden layers (the 1^st^, 2^nd^, 3^rd^, and 4^th^ hidden layer, in the direction of forward propagation), which had 128, 64, 16, and 8 nodes, respectively. The 1^st^, 2^nd^, and 3^rd^ hidden layers were each followed by leaky ReLU activation layers (negative slope 0.01), dropout regularization layers (dropout probability 0.4, 0.4, and 0.2, respectively), and batch normalization layers. The fourth hidden layer was followed by a sigmoid activation layer. Starting learning rate was 0.0001. Binary cross entropy loss function and cosine annealing learning rate scheduler were used for model optimization.

For the convolutional neural network, models were trained using the PyTorch framework, in mini-batches of size 128. For the best performing model, the neural network consisted of 4 hidden convolutional layers and 2 hidden fully connected layers. The first convolutional layer (6 filters; filter size:1; stride:1) was followed by a batch normalization layer. Other convolutional layers (6 filters; filter size:4; stride:4) were each followed by batch normalization layers, leaky ReLU activation layers (negative slope 0.01), and max pooling layers (filter size:2; stride:2), The first fully connected layer (16 nodes) was followed by a leaky ReLU activation layer (negative slope 0.01) and a batch normalization layer. The second fully connected layer was followed by a sigmoid activation layer. Starting learning rate was 0.001. Binary cross entropy loss function and cosine annealing learning rate scheduler were used for model optimization.

Random Forest and AdaBoost classifiers were trained with the scikit-learn (version 1.0.2) library (Pedregosa et al., 2012). Grid searches were performed via 5-fold cross-validation to optimize the accuracy. The best performing Random Forest classifier had 500 estimators and a maximum depth of 2. The best performing AdaBoost classifier had 300 estimators and was trained with a learning rate of 0.1.

### Model Interpretation with SHapley Additive exPlanations (SHAP)

SHAP analysis was performed with the DeepExplainer function of the SHAP library (version 0.40.0) (Lundberg and Lee, 2017). We generated three models with similar performances, using different seeds for random splitting of the datasets. Based on our computational resources, we used these models to compute SHAP values of 1000 cells randomly selected from the whole dataset. SHAP values used for biological interpretation were computed by averaging the SHAP values derived from the three models. For cell subtype-specific SHAP analysis, we computed SHAP values of 1000 randomly selected cells corresponding to each cell subtype, or all cells of that subtype if their total number was less than 1000.

### Pathway Enrichment Analysis and Protein Interaction Network Analysis

Gene Ontology pathway enrichment analysis was performed in the online portal of ShinyGo version 0.76 (http://bioinformatics.sdstate.edu/go/) (Ge et al., 2020). Genes with the top 10% (90^th^ percentile) median absolute SHAP values were compared to all genes in the dataset (background genes; top 10% genes included). False discovery rate cut-off was set to 0.05.

Protein interaction network analysis was performed in the online portal of STRING version 11.5 (https://string-db.org/) (Szklarczyk et al., 2019). Genes with the top 1% (99^th^ percentile) median absolute SHAP values were analyzed using the “Multiple proteins” entry. To select only the high confidence interactions, the minimum required interaction score was set to 0.7.

### Differential Gene Expression Analysis

Differential gene expression analysis was performed with the FindMarkers function of Seurat. By default, absolute log2(fold change) threshold was 0.25 and Wilcoxon Ranked Sum test was used to determine *p* values.

## Supplemental Excel Tables

Table S1. Ranked median SHAP values of all genes.

Table S2. Ranked cell subtype-specific median SHAP values of all genes.

Table S3. Differentially expressed genes between PHA-L^high^ and PHA-L^low^ cells in the TIL dataset.

