## Supplemental Figures for "Deep Learning Explains the Biology of Branched Glycans from Single-Cell Sequencing Data"

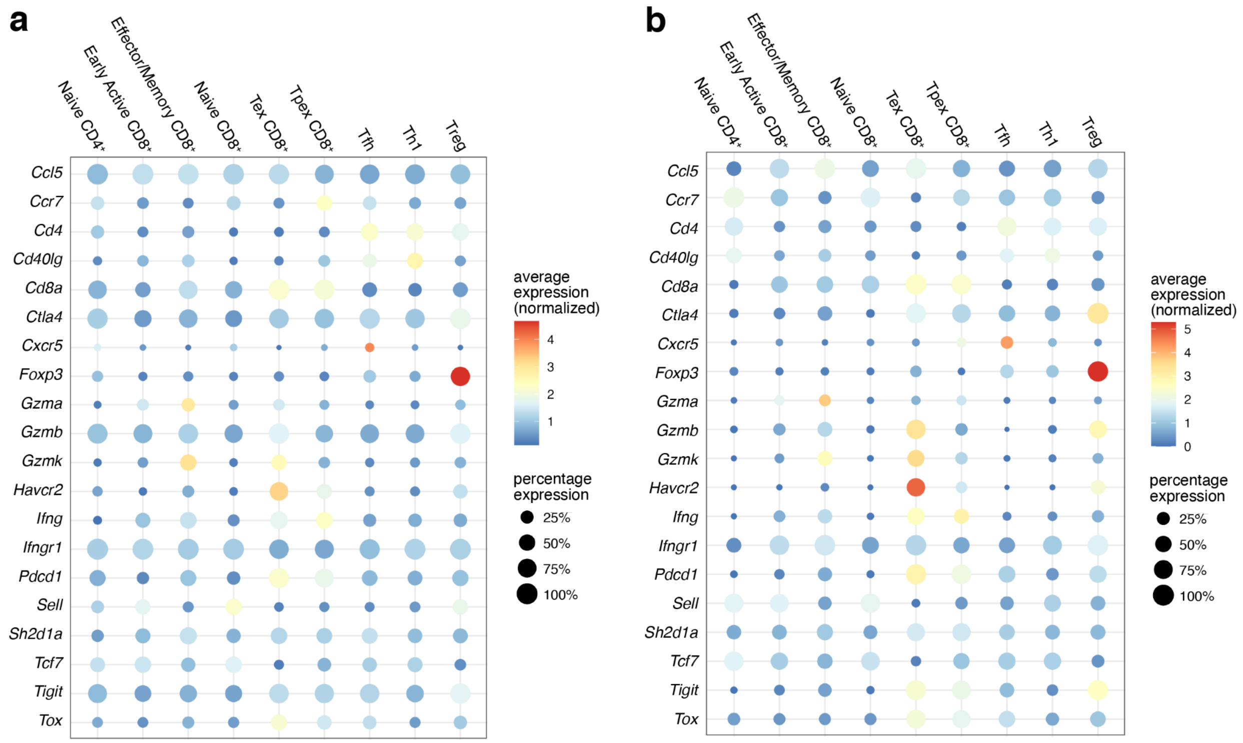


**Figure S1.** Expression of marker genes of subtypes of T cells in the (**a**) TIL dataset and (**b**) LN dataset. For each gene, the average expression has been normalized to the mean of expression of the gene in all types of cells.

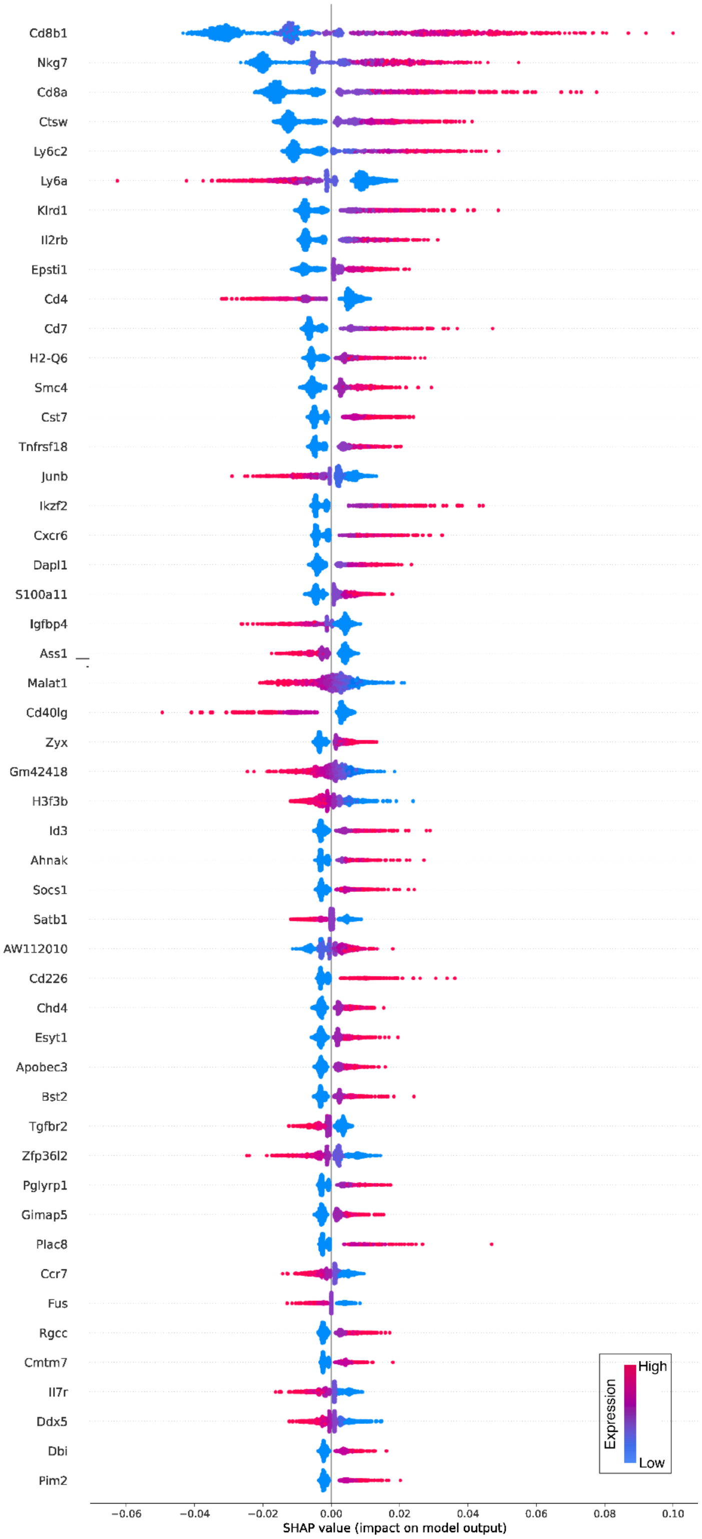


**Figure S2.** SHAP values of top 50 genes in the LN dataset ranked by median absolute SHAP value.

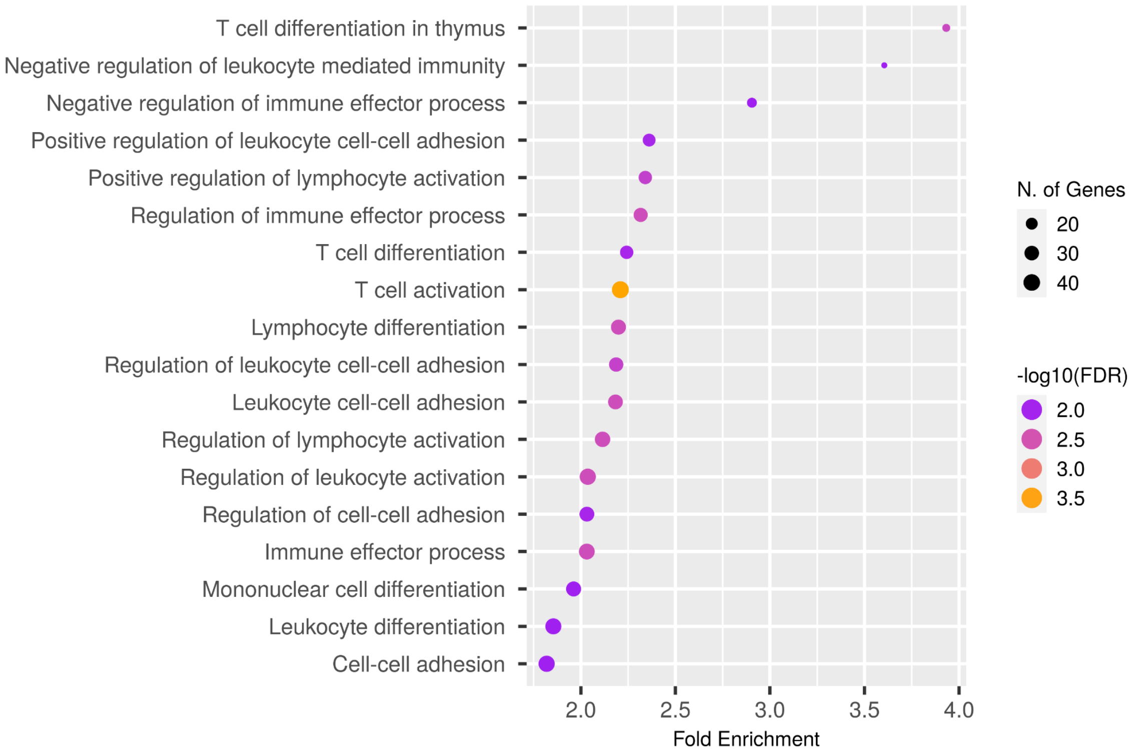


**Figure S3.** Gene Ontology pathway enrichment analysis of using the SHAP genes of the LN dataset.

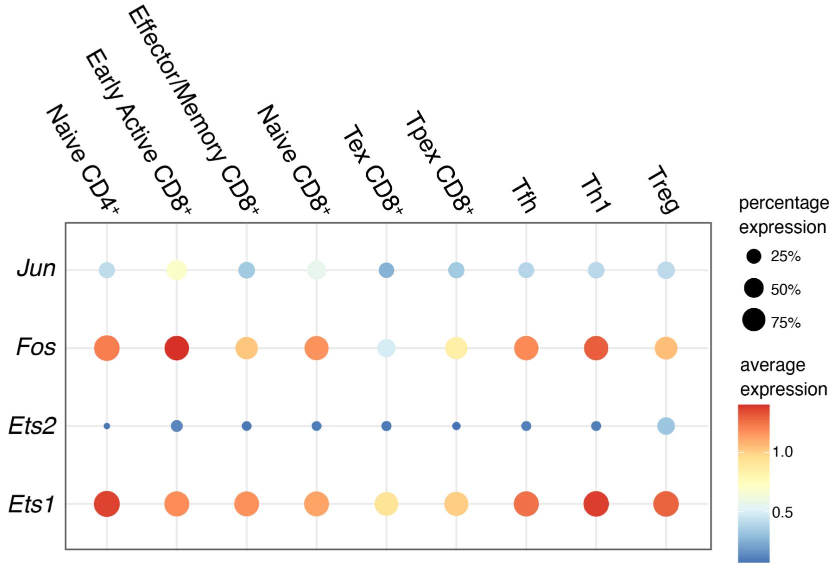


**Figure S4.** Expression of four genes encoding transcription factors regulating MGAT5 expression across subtypes of T cells in the TIL dataset.
